# Efficiency of island homing by sea turtles under multimodal navigating strategies

**DOI:** 10.1101/453431

**Authors:** K. J. Painter, A. Z. Plochocka

## Abstract

A dot in the vastness of the Atlantic, Ascension Island remains a lifelong goal for the green sea turtles that hatched there, returning as adults every three or four years to nest. This navigating puzzle was brought to the scientific community’s attention by Charles Darwin and remains a topic of considerable speculation. Various cues have been suggested, with orientation to geomagnetic field elements and following odour plumes to their island source among the most compelling. Via a comprehensive *in silico* investigation we test the hypothesis that multimodal cue following, in which turtles utilise multiple guidance cues, is the most effective strategy. Specifically, we combine agent-based and continuous-level modelling to simulate displaced virtual turtles as they attempt to return to the island. Our analysis shows how population homing efficiency improves as the number of utilised cues is increased, even under “extreme” scenarios where the overall strength of navigating information decreases. Beyond the paradigm case of green turtles returning to Ascension Island, we believe this could commonly apply throughout animal navigation.

## 1. Introduction

Sea turtles are expert navigators, capable of impressive transoceanic migrations from hatchling to adult [34, 36]. Highlighted by Darwin [10], the Ascension Island (AI) green turtles return to their secluded island nesting beaches during triennial return journeys that span the thousands of kilometres of open ocean from foraging grounds along the South American coast. Ideally turtles could be tracked between Brazil and AI and back, yet this remains infeasible with di culties ranging from locating pre-migratory turtles in the open ocean to equipment limitations. Conveniently, females nest repeatedly [41, 64]: finding and displacing a nesting turtle is (relatively) easy, and their rehoming attempt can subsequently be tracked. Even then challenges remain in terms of untangling the data from uncertainties, such as the variable currents and an individual’s necessity to return. Laboratory studies allow even greater control, yet these are limited to hatchlings or juveniles and, inevitably, questions remain on extrapolating to the real world. Beyond experiments, theoretical modelling permits total control of the underlying inputs, although this is also its failing in that a model is only as good as its assumptions. Nevertheless, modelling o ers a counterpoint to experiment and can be used to test theories for navigation[29, 53, 51, 56, 21, 46, 14, 58].

To achieve migration, animals are commonly assumed to detect various cues to formulate a “map and compass”, where the map provides position relative to a target and the compass gives a heading [49, 43]. Of the potential cues, several can be discounted to some degree: sun/light could provide coarse-level navigation [42], yet poor (outside water) vision would probably exclude precise celestial navigation [11]; ocean depth and land absence would also discount topographic orientation, at least until close. Two leading theories that persist, however, are based on responses to the geomagnetic field and following an odour plume to its island source.

Numerous laboratory studies implicate geomagnetic field responses: hatchlings display distinct swimming orientations under artificial fields that vary in either magnetic inclination angle [32] or intensity [31]; investigations with hatchling [30] and juveniles [33] show subtle changes to their preferred heading for the di erent fields encountered along their migratory routes. Extracting latitude from the magnetic field is certainly conceivable, given how the geomagnetic field changes with latitude, yet longitude can also be inferred [50] and therefore magnetic field sensing can o er bi-coordinate positioning. Recent field studies using displaced adults provide further support [38, 5], where turtles perturbed by artificial magnetic fields show impaired homing. Notably total intensity and inclination angle isoclines lie almost orthogonally (see Fig. 1) at AI, facilitating its identification via these field elements.

**Figure 1:**
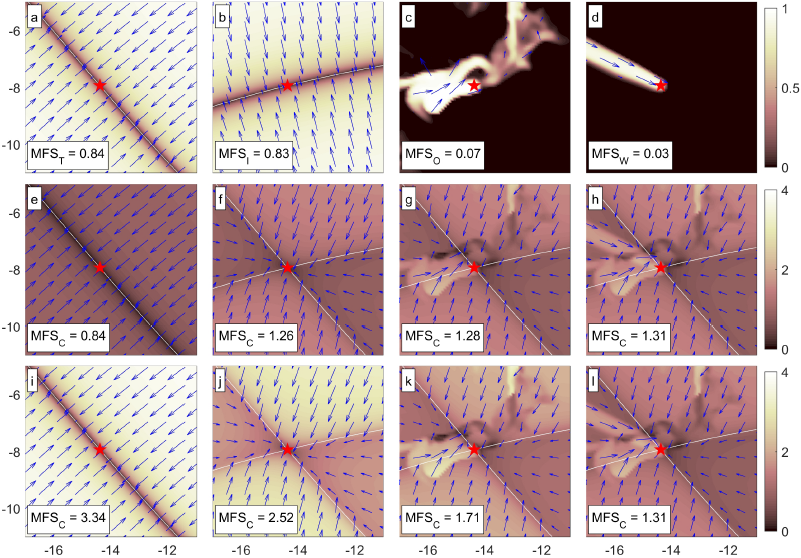
Individual and consolidated navigating fields. Arrows indicate the preferred direction, with their length and the colour density map indicating the strength/certainty. AI indicated by central star. (a-d) Individual navigation fields: (a) w_T_; (b) w_I_; (c) w_O_; (d) w_W_. (e-l) Consolidated navigation fields, w_C_ =: (e,i) w_T_; (f,j) w_T_ + w_I_; (g,k) w_T_ + w_I_ + w_O_; (h,l) w_T_ + w_I_ + w_O_ + w_W_. (e-h) and (i-l) respectively show pathways of cue addition under accumulative (*c* = 1) and redistributive normalisation (*K* = 4), leading to the same field when all cues are used (h,l). Snapshot as of 15/02/14; a movie is included in the supplementary information (SM1).

The idea of navigation in response to island-generated odour plumes was introduced in [29]. The South Equatorial current passes near the island, yielding a general east to west flow, while trade winds are persistently from the south east. Hence, either mechanism could generate a reasonably consistent plume. The study of [29] showed ocean-transported compounds (under plausible production, convection and di usion rates) could form a detectable plume hundreds of kilometres away. Laboratory studies reveal numerous air/waterborne substances that elicit responses, varying from specific molecules to coastal mud [39, 40, 18, 15, 12, 13]. Displaced turtles also appear to find it easier to return to AI from the north-west than south-east [20], consistent with the direction of a wind-transported plume.

Caveats must be attached to either theory. Magnetic di erences diminish closer to the source, so exact pinpointing would demand exquisite sensitivity. Yet this would render turtles susceptible to spatial and temporal fluctuations in the field: spatial anomalies occur with changes in the local geology while temporal fluctuations vary with solar activity. Over longer time-scales, secular change is considerable over the three year or so interval between migrations. Theoretical considerations into the impact of secular variation on a magnetically-guided population are given in [52, 35], while focussed studies have specifically examined its relationship with the shifting pattern of loggerhead sea turtle nest sites [9]. Given these issues, magnetic sensing is perhaps more likely to lead a turtle to general island proximity than allow its precise localisation [1, 34, 5, 43]. Regarding plumes, beyond which precise substance(s) are followed, questions form on its spatial extent. Even under highly refined senses, prevailing wind/currents will leave large regions untouched by the plume to create blind spots. Further, to provide guidance, detection must be coupled to upflow movement: anemotaxis for oriented movement to air currents and rheotaxis for water currents. Rheotaxis for an individual immersed in water and far from landmarks is far from trivial and direct evidence in turtles has proven somewhat di cult to obtain, although some recent analysis o ers support [28].

The strengths and weaknesses associated with the hypotheses above have led several to consider a multimodal/combinatorial navigation strategy, in that turtles integrate the directions suggested by di erent cues in a manner that allows them to robustly pinpoint their destination: for example, see [1, 34, 5, 14]. Thus, as one potential example, magnetic field information could lead them to the general vicinity of the island before a plume is encountered and followed upwind/upcurrent. A recent theoretical study of [14] examined the intersection between magnetic field elements and transported odour plumes about AI, concluding that a multimodal strategy whereby both are utilised could prove more effective. Similarly, agent-based simulations in [58], albeit within a more abstract setting, have investigated the usefulness of multimodal strategies based on magnetic guidance and plume following. Consistent with these theories of multimodal based homing, magnetically-disturbed turtles that show impaired homing can still eventually home, suggesting that other cues are utilised when an information source is removed [47, 38, 5].

In this paper we perform a formal theoretical test into the extent to which multimodal homing proves more e cient navigation. A hybrid agent-based/continuous (ABM) and its corresponding fully-continuous model (FCM), obtained through scaling, are used to track virtual turtle paths/population distributions in an *in silico* displacement study. An evolving navigating field o ers guidance, derived from geomagnetic field data and computed current or wind transported odour plumes. Utilising more cues is shown to substantially improve the rehoming success rate of the AI turtle population.

## 2. Methods

### Study region and release dates/locations

The *in silico* study displaces virtual turtles from the island and tracks their rehoming over a period of up to *t*_end_ = 100 days. Our study region centres on x_*AI*_ = 14.35°W, 7.94°S (the approximate centre of AI) and stretches 10° s in latitudinal and longitudinal directions. *Home* is a circular area of radius 15 km and centred on x_*AI*_, describing a region that extends ~10 km from the coastline. Once in this range, we assume visual, auditory and other senses bring the turtle to AI. Note that tracked turtles have approached as close as 23 km without returning [37]. Turtles are assumed to spend the majority of time at or near the surface, allowing us to restrict movements to a two-dimensional plane. Of course, this is a simplification, as navigating green turtles can perform deeper dives [19] and could potentially use these to locate more deeply-located cues or limit their exposure to strong currents.

Variable ocean currents substantially alter the ability and/or time needed to home [46], and we reduce their impact as follows. First, conditions at a specific release location (e.g. a point NE) are limited by displacing each turtle in a population (of size *N* = 100) to a point randomly located along a circular corridor surrounding the island (mean initial distance from AI = 300 km). Second, the effect of temporal changes are reduced by averaging over multiple releases spanning multiple years, with populations released on 1st February/April for each of the years 2010-2015 (the middle of the nesting season).

Homing success is measured by the *homing efficiency*,

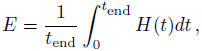

swhere *H*(*t*) is the population fraction returned home by time *t*. Values of *E* close to 1 define effective navigators that return quickly following release, while values close to 0 represent poor navigators that require a long time to home, if ever.

### Simulation methods

Virtual tracks and population distributions are computed by adapting a previously developed multiscale framework [46]. In the ABM virtual turtle agents are immersed into a flow field representing ocean currents. The FCM describes the corresponding non-homed population density distribution *p*(x, *t*) at position x and time *t*. Here we outline the key points with details provided in the Appendices. Briefly, each virtual turtle orients according to a consolidated navigation field (C) generated from responses to up to four individual cues: magnetic total intensity (T); magnetic inclination angle (I); an ocean-borne chemical plume (O) and a wind-borne chemical plume (W).

Agent motion in the ABM derives from ocean current drift and oriented swimming, the latter described by a “velocity-jump” random walk [45] consisting of smooth swims with fixed heading interspersed by reorientation. Note that detailed tracks obtained from “crittercams” attached to loggerhead turtles suggest this to be a reasonable approximation of oriented swimming behaviour [44]. Further to vector fields for ocean currents (uocean), the critical statistical inputs for parametrising this model are: (i) the average speed (*s*); (ii) the average rate of reorientations (λ); (iii) a directional distribution *q*(α) for turning into a new heading, angle α.

The distribution *q* allows navigation to be entered, via biasing turns into specific angles. We employ the von Mises distribution, a standard in animal navigation studies [4]. Specifically,

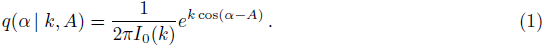

The parameters *k* and *A* denote the *navigational strength* and *dominant angle* respectively: large *k* generate a majority of turns within a few degrees of the dominant angle *A*, while for negligible *k* no particular angle is favoured. Cues typically vary spatially and/or temporally, so *k* and *A* are functions of *t* and x. *I_j_*(*k*) denotes the modified Bessel function of first kind (order *j*) and enters as a normalising factor.

The FCM for *p*(x, *t*) is a partial di erential equation of advection-anisotropic di usion type, obtained through scaling the ABM. Hence, its terms and parameters directly follow from the statistical inputs to the ABM (see Appendix A and [46]). The advantages of this dual modelling approach are: (i) the ABM is formulated at the level of individual movement, facilitating parametrisation against typical tracking data; (ii) the FCM is computationally cheap and tractable, allowing broader (less parameter-specific) analyses.

### Navigating fields

The distribution (1) is parametrised via the *consolidated navigation field*, a vector field w_C_(x, *t*) that combines individual cue fields into a single entity that confers navigation information. The individual cue fields are taken to be as follows:

- w_T_ and w_I_, respectively describing responses to geomagnetic total intensity and inclination angle;
- w_W_ and w_O_, respectively describing responses to windborne and oceanborne odours.

Each of these are characterised by two parameters, *k_i_* and *_i_*, where *i* ∈ T,I,O,W. The former is a strength measure for the maximum “certainty” by which an agent follows a particular field, while the latter reflects the sensitivity of detection.

Detailed equations are provided in Appendix B and here we restrict to the key ideas informing the choices. Navigation to intensity and/or inclination assumes an innate ability to detect and recall their values. Specifically, we assume these are imprinted while at the island (see [35] for a theoretical discussion) and, following displacement, orientation biases are experienced according to the di erence between the current location/island values. Temporal variation (diurnal, secular) and spatially localised anomalies in the field are ignored and the intensity/inclination inputs are given by the mean values in a standard geomagnetic field model. We should emphasise, however, that this is a simplification and future modelling will extend to account for fluctuations of the field, see discussion for further comment. Typical fields for w_T_ and w_I_ are illustrated in Fig. 1 (a-b). Navigation to single field elements orient individuals to an isocline intersecting the island, with response strength increasing with distance from it.

Responses to odour plumes tend to follow a general pattern of detection leading to movement upflow [61]. We assume the strength of response increases (and saturates) with concentration, with subsequent movement against the current/wind direction. This demands an additional equation to describe the odour concentration dynamics, *c*(x, *t*), taken to be of advection-di usion-reaction type (see Appendix B). Key inputs are velocity fields to describe transport by ocean currents (uocean) or surface winds (u_wind_), a di usion coe cient *D*, a source term characterised by the substance production rate *M* and a decay term measured by the half life *τ*. Snapshots of instantaneous navigation fields generated by an ocean or wind plume are illustrated in Fig. 1 (c-d). We note the considerably larger, yet more contorted, nature of the ocean plume generated field.

Individual fields are combined into the consolidated field through simple summation (w_C_ = w_T_ + w_I_ + w_O_ + w_W_). We subsequently take *k*(x, *t*) = *|*w_C_(x, *t*)| and *A*(x, *t*) as the angle in the direction of w_C_(x, *t*) in Equation (1). In words, an individual navigates in the local direction of w_C_(x, *t*) with certainty determined by its local length. Consolidated field plots are illustrated in Fig. 1 (e-l) based on the individual fields in (a-d).

Note that 0 ≤ *k*(x, *t*) ≤ *k*_T_ + *k*_I_ + *k*_W_ + *k*_O_ = *K*, where we call *K* the *maximum navigating strength*. This strictly theoretical maximum only applies if all individual fields point in the same direction at their maximum strengths, yet its concept provides a reference for normalisation. We define the mean field strength and mean field utilisation, respectively

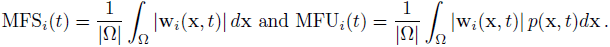

The former is a general measure of a cue’s strength across the entire study region (Ω), while the latter indicates the extent to which it is utilised. Note that w_O_ and w_W_ yield considerably lower mean field strengths (Fig. 1 (c-d)), reflecting how plumes “contact” only a fraction of the study region at any given instant.

### Datasets, parameters and normalisation

Ocean currents, surface winds and magnetic field components are obtained from standard (public domain) datasets: HYCOM for ocean currents [8], ASCAT measurements for surface winds [6] and the IGRF model [60] for the magnetic field, see Appendix D.1. Model parameters utilise the default parameter set in Table D.2. Where possible, these are estimated from data and references are provided in Appendix D.2.

Multimodal combinations are implemented through distinct choices for (*k*_T_, *k*_I_, *k*_O_, *k*_W_). Setting a particular *k_i_* to zero eliminates the corresponding cue from the orientating response. We set *n* as the number of utilised cues and, to compare across combinations, choose two normalisations:

- In *accumulative* normalisation adding a cue does not alter the response to other cues, it simply adds the new field. Specifically we take *k*_i_ *2 {*0, *c}* and hence *K* = *nc*.
- In *redistributive* normalisation strategies are compared at fixed values of *K*, equally shared across contributing cues: *k*_i_ *2 {*0, *K/n}*, so adding a new cue is countered by diminished responses to existing cues.

These two forms are demonstrated in sequences Figure 1(e-h) and (i-l). Accumulative normalisation typically yields a rise in MFS_C_ as cues are added. Redistributive normalisation, on the other hand, generally decreases it. In this sense, these two methods can be considered as extreme scenarios.

## 3. Results

### Single release date

We begin the investigation using a release date of 01/02/2014 and show simulations of the ABM and FCM under selected strategies. It is noted that FCM output quantitatively describes that of the ABM, and hence o ers meaningful data on average population behaviour. Fig. 2(a) combines plots of agent positions (ABM) with the continuous population distribution (FCM) at various times following release. In this simulation, navigation is based on intensity and inclination alone: the bicoordinate information ensures the dominant direction is towards home from any point, but with a certainty diminishing with proximity.

**Figure 2:**
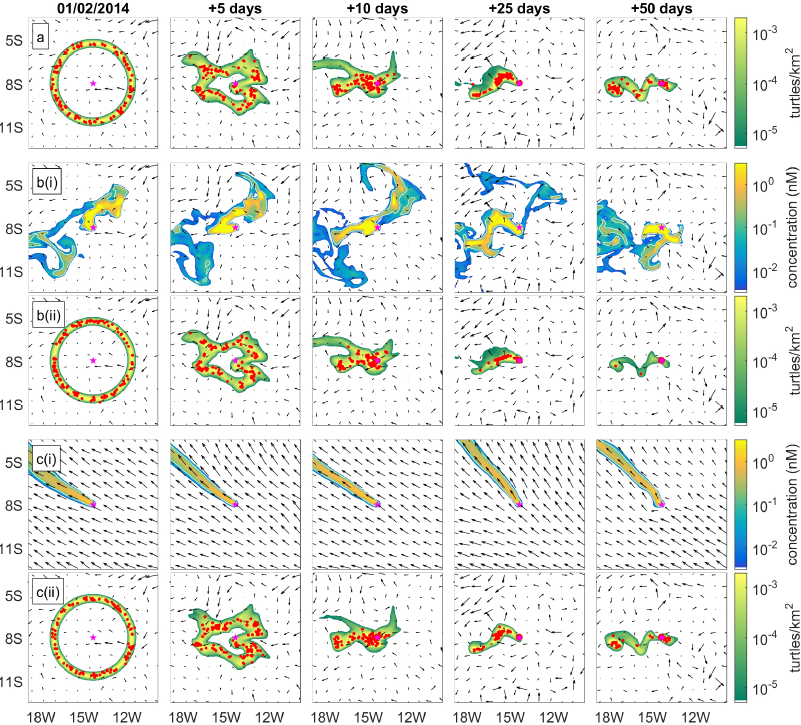
Simulations of turtle homing for (*k*_T_, *k*_I_, *k*_O_, *k*_W_) = (a) (1, 1, 0, 0), (b) (1, 1, 1, 0), (c) (1, 1, 0, 1). In a, b(ii), c(ii) turtle positions (red circles) from the ABM overly *p*(x, *t*) (colourscale) computed from the FCM. Arrows indicate direction and strength of the ocean currents. In b(i) oceanborne and c(i) windborne plume concentrations (colourscale) are plotted, overlaid with arrows indicating currents and wind velocities. All populations released on 01/02/2014, parameters as in Table D.2. Movies for (b,c) are included in the supplementary information (SM2, SM3).

The population of Fig. 2b(ii) is given an additional sense of ocean plume navigation. Turbulent currents fashion a contorted and meandering plume (Fig. 2b(i)) that occasionally generates contradictory information (pointing away from AI), but also ensures much of the field “encounters” it at some point: an average of only 6% of the study region experiences concentrations exceeding the plume sensitivity threshold at a particular instant, but a cumulative 38% is covered over the full simulation (Fig. 3 (a)). The overall contribution is positive and, for this population at least, turtles take quicker return paths. Fig. 2c shows the corresponding case where the population is imbued with additional navigation to a windborne cue. Persistent winds create a relatively stable plume that shifts little: instantaneous and cumulative encounter fractions are just 4% and 20% (Fig. 3 (b)). On the other hand, the pull is more consistently towards home and the extra capacity again hastens turtle homing times.

**Figure 3:**
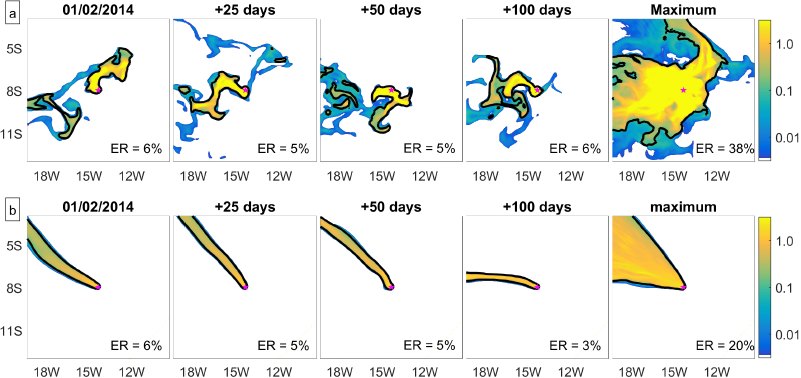
Figures showing the spatial extent of (a) oceanborne and (b) windborne plumes across the study region. The first four columns show instantaneous plumes on the given days, while the final column plots the maximum concentration experienced across the full 100 days of the simulation study. The colourscale indicates the concentration, expressed in *nM*. The (instantaneous or cumulative) encounter region (ER) is the percentage of the study field that experiences a concentration above the corresponding sensitivity parameter (κ_O_ or κ_W_), with the region of the field meeting this criterion indicated in the plots as the area enclosed by the solid black line.

Population distributions have similar shape for each strategy: they only di er in the addition or not of odour-based responses, so only a fraction lying inside a plume behaves di erently. Nevertheless, these small di erences can lead to sizeable increases in the homed population, Fig. 4(a-c). To understand this we examine the mean field strength (MFS_i_) and mean field utilisation (MFU_i_), Fig. 4 (d-f). The mean field strength of the consolidated field changes minimally with each scenario, with magnetic field elements dominating the contribution. Plume importance, however, is reflected in the individual cue utilisation: spikes of significant ocean/windborne cue utilisation appear with increases in the homed population. E ectively, these cues can help nudge turtles home once the magnetic field has bought them su ciently close.

**Figure 4:**
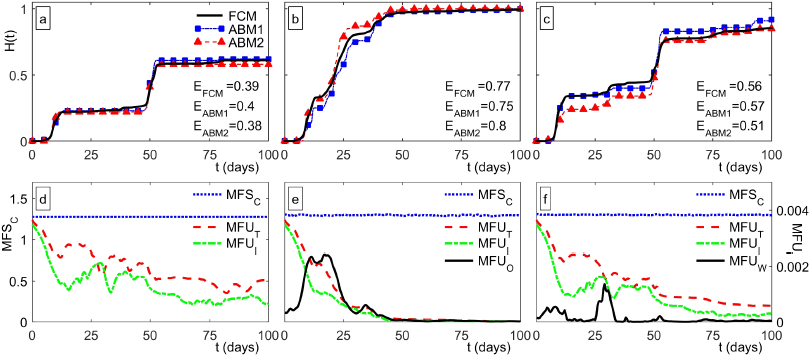
Summary data for simulations in Figure 2. (a-c) *H*(*t*) and *E*; key in (a) indicates results from the FCM and two separate realisations of the ABM. (d-f) MFS_C_(*t*) (left-scale in (d)) and MFU_i_(*t*) (right-scale in (f)). (a,d) correspond to Fig. 2(a); (b,e) to Fig. 2(b); (c,f) to Fig. 2(c).

### Efficiency of multimodal strategies

Homing is assessed under a full range of multimodal strategies. For the available cues we have 16 distinct combinations, ranging from utilising none to all 4. Comparison is made following additive normalisation (AN) and redistributive normalisation (RN). The initial exploration utilises the ABM, where we release 100 turtles on 01/02/2014 for each strategy (Table 1). A cursory glance suggests utilising more cues generally improves homing, with individuals travelling shorter distances and adopting straighter paths.

**Table 1:**
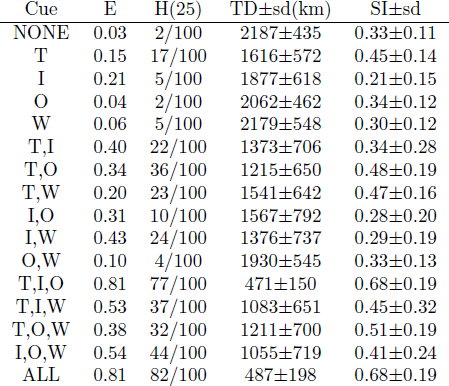
ABM data for turtle homing attempts, showing cue combination, *E*, *H*(25), mean track distance (TD), mean straightness index (SI); see Appendix D.3 for formulae. In each case, *N* = 100 turtles are released on 01/02/2014; AN is applied with *c* = 1.

We systematically investigate this in a large-scale analysis, exploiting the computational convenience of the FCM and minimising current bias by averaging over all 12 release dates. Results of the analysis are reported in Fig. 5 for the sixteen combinations under (a-c) AN and (d-f) RN. We plot both *E* and a more conventional measure, the fraction homed after 25 days. The leftmost bars in (a,d) correspond to a complete absence of cues (random movement): unsurprisingly, this yields the lowest efficiency and gives a baseline for comparison.

**Figure 5:**
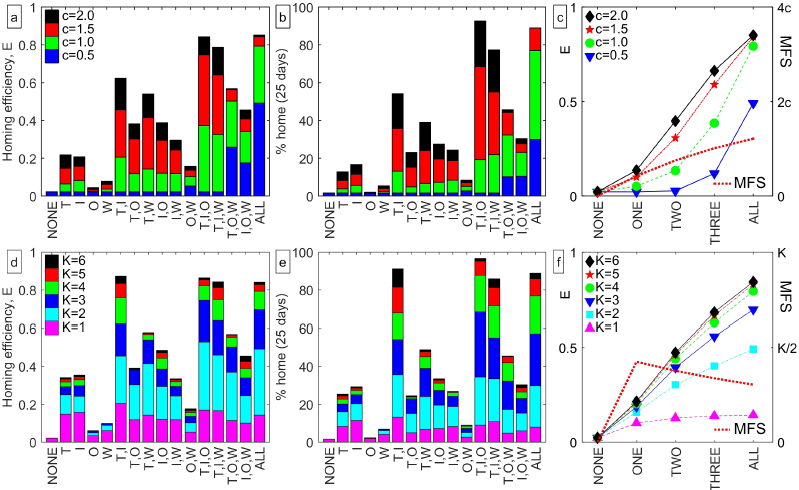
Homing efficiency of multimodal strategies under (a-c) AN and (d-f) RN. (a,d) *E* and (b,e) *H*(25) for each combination. Colours indicate how *E* changes with field strengths (a,b) *c* or (d,e) *K*. (c,f) Mean value of (left axis) *E* and (right-axis) MFS, when averaged across strategies using none to all of the available cues.

The analysis supports the notion that more cues leads to more e cient homing (Fig. 5 (c,f)), even under the extreme redistributive normalisation. Included in these plots are the mean navigating field strengths, averaged over time and cue combinations and plotted as a function of the number of cues. Average homing efficiency increases as a function of the number of cues utilised, even when the mean field strength decreases (under RN). Homing efficiency increases with field strength parameters but eventually saturates: natural constraints lie in the swimming speed and the general ocean currents, even for the most precise homing. Year by year analysis shows fluctuations to the efficiency of a particular strategy according to release date, Fig. 6 (a,c), yet the overall trend is consistent and, averaged across all strategies, homing efficiency changes little with release date, Fig. 6 (b,d).

**Figure 6:**
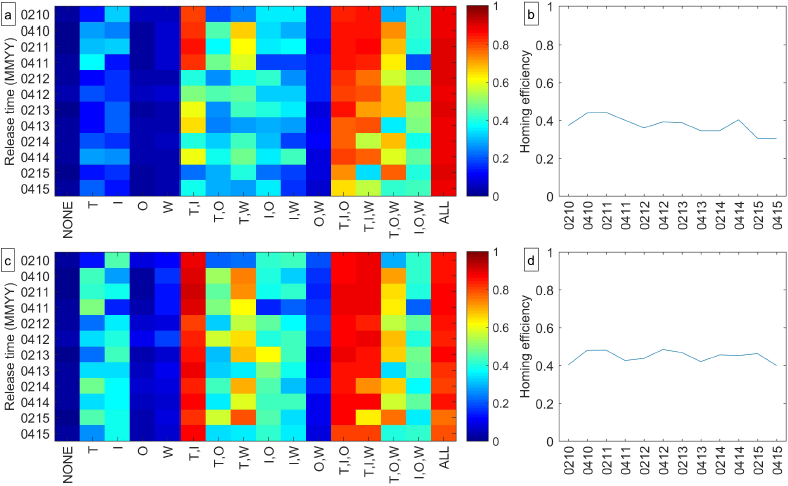
(a,c) Homing efficiency plotted as a function of release date and strategy. (b,d) Homing efficiency averaged over all combinations and plotted as a function of release date. (a,b) use accumulative normalisation with *c* = 1 while (c,d) use redistributive normalisation with *K* = 4.

Single cue strategies generate low homing e ciencies, even under high navigational strengths: odour-based strategies demand finding and remaining in the plume; orientation to single field properties allow movement to an isocline bisecting the island, but not exact pinpointing. Dual-combination strategies are considerably more effective with, in particular, a response to intensity and inclination generating e cient homing: this is perhaps unsurprising given its capacity to provide bicoordinate information. Of other dual-cue strategies, a combination of intensity and windborne cues is more effective than others. This combination benefits from intersection between the intensity isocline and the wind-generated chemical plume (see also [14]): turtles navigate to the isocline and locate the plume.

To evaluate cue addition more precisely we construct network graphs (Fig. 7) where lines connect di erent permutations from none to all four cues. Each connecting line illustrates the positive/negative impact on *E*: AN generates uniformly positive effects, Fig. 7(a); under RN additions are overwhelmingly positive (25/32), Fig. 7(b). Surprisingly, cue additions can generate highly positive effects on homing efficiency, even when there is a significant drop in the mean navigating field strength (compare black lines in Fig. 7 (b) with red lines in (d)). Moreover, where any negative connections do occur they lead to only a marginal decrease in the homing efficiency despite a correspondingly large decrease in the mean navigating strength (see Fig. 7 (d)). In other words, the use of multimodal sensing can overcome significantly reduced field strength and allow individuals to home almost as e ciently. Similar trends can be observed at di erent parameter sets (data not shown). Overall, the results strongly support the notion that multimodal homing strategies are more effective.

**Figure 7:**
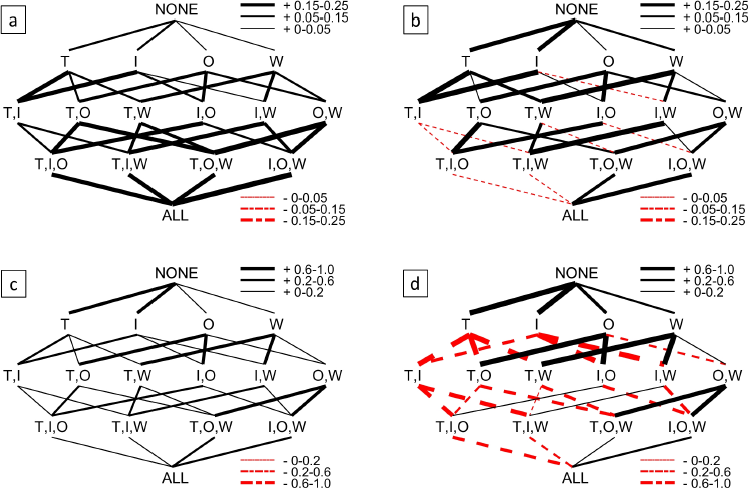
(a-b) Networks illustrating effect of systematic cue addition on homing efficiency: (a) AN and (b) RN. (c-d) Networks illustrating effect of systematic cue addition on mean navigating field strength MFS_*C*_: (c) AN and (d) RN. In all plots, solid black (dashed red) lines show increased (decreased) homing efficiency or MFS_*C*_, with the size of the change represented by line thickness. Changes are obtained by averaging over the 12 release dates and over the full simulation time course.

## 4. Discussion

Intuitively, relying on a single navigating cue, however potent, would render a population sensitive to natural or otherwise change. For species such as sea turtles, whose survival depends on their ability to find specific nesting beaches, robust population-level navigation is crucial. Logically a multimodal strategy should prove more effective and we provide a formal theoretical test of this in an exemplar of animal navigation, the homing of Ascension Island green turtles. The investigation reveals that multimodal strategies improve homing, even following strict normalisation: specific simulations show how the additional boost from ocean and/or windborne cues can nudge magnetically-guided turtles home.

When single strategies are employed, magnetic appears to outperform chemical quite substantially. Directly extrapolating this to the AI case, though, must be approached cautiously: the need for a tractable model demands simplifications which can over or understate di erent factors. Particularly, it is likely that our model underestimates the potential guidance of chemical plumes. Following an odour demands finding and remaining inside a complex plume: in the absence of the cue, our agents perform a naive unbiased random walk (Brownian motion), a logical abstraction yet one that probably oversimplifies more e cient searching behaviours, such as a Levy walk (e.g. [54]). Similarly, more sophisticated plume-following such as “zigzagging” [61] may reduce the likelihood of losing contact with a plume when found. On the other hand, we can imagine the contribution of magnetic cues to be overemphasised: smooth and fixed magnetic field data o ers reliable guidance, yet natural magnetic fields are considerably rougher and noisier with diurnal variations alone in the 10-50 nT range, similar to chosen sensitivity parameters. While such fluctuations are implicitly accounted for in the strength parameters (a measure of maximum cue certainty), a future study may benefit from explicitly incorporating variation in field values, allowing exploration into whether they reduce reliability (fluctuations misleading turtles), can benefit homing (e.g. following local field anomalies) or how turtles may minimise the impact. Theoretical investigations, using agent-based modelling, addressing similar questions have been initiated in [59]. Over longer periods secular variation becomes significant: in the 3/4 year inter-migration period, magnetic positions can change 50–100 km in the region of AI. The naive agents here would struggle under such change, and how they meet this challenge is an important issue to address. Overall, however, regardless of whether certain cues are over or underestimated, none of the single-response strategies appeared particularly effective.

Our focus explicitly explored mid-range (10-100s of kilometres) journeys, rather than short-range (close to the island) or long-range migration from the South American coast. Investigating the former would involve reconsidering our home definition: a circular region ensures no approach angle is favoured, but the distance of confident identification is likely to vary about AI; for example, wave refraction patterns, sound propagation and benthic topography will all vary about the island. At longer distances additional cues such as a relatively simple sun-compass may also factor into the navigation [42]. Nevertheless, given the di culty of field studies, modelling the full migration path from South America to AI forms a key objective for future work. Cues are believed to provide effective navigation across specific ranges – for example, in the case of a magnetic map the resolution is assumed to be down to the order of 10 − 50 m [43] – and therefore a detailed study may shed light on which cues are utilised during di erent migration stages.

Multiple cues were simplistically combined through linear addition, facilitating automation. Other methods may investigate a more nuanced evaluation, such as cue prioritisation (choosing one over another if perceived to be more reliable). Implicit in our assumptions (for oceanborne cues) is an ability to perform rheotaxis. This capacity is far from trivial: for turtles it could potentially arise from passive shell twisting from flow dynamics, inference from surface wave direction or detection of water flow along specialised sensory organs. Rheotactic responses are documented for various species and recent analyses for turtles provides some support [28], if not definitive proof. Without rheotaxis, it is not clear how an ocean plume could o er clear navigating information, although possible chemical cue dectection could generate a simpler kinesis-type response. If rheotaxis were possible, it is also tempting to consider a potentially wider influence such as directional movement changes/deeper diving to avoid disadvantageous currents.

A flip side to our findings is, of course, that homing will be impaired under perturbation to the underlying cues. Slower island homing times are observed for Indian Ocean green turtles subjected to carefully positioned magnets [38, 5]. Ocean currents, winds and magnetic field properties (as well as the potential availability of other navigating cues) will all be distinct from those surrounding AI, and an important extension would be to investigate the generality of our results within that population. Navigating cues can potentially be disrupted through natural or otherwise change: magnetic fields are perturbed following seismic activity or during solar storms; the persistent presence of chemicals following an oilspill could reduce the capacity of turtles to detect other substances. The subsequent impact on homing capability, and subsequently population-level nesting, could form a test for future studies.

## Appendix A. Mathematical Models

### Appendix A.1. Hybrid Agent Based Model (ABM)

In the ABM each agent is defined by a point Lagrangian particle that moves continuously in a flowing medium. Agent motion derives from locally-encountered ocean currents (passive drift) and oriented swimming (active motion). The currents are described by a velocity vector field uocean(x, *t*) (x is position and *t* is time) obtained from public-domain datasets. For simplicity, turtles are assumed to remain at roughly the same depth and we can therefore restrict to a two-dimensional field. Oriented swimming is described by a “velocity-jump” biased random walk [45], in which movement occurs as smooth runs with constant velocity, punctuated by (e ectively) instantaneous reorientations into new headings. Detailed tracks from loggerhead turtles attached with “crittercams” suggest this to be a reasonable approximation of real-life patterns, where directed swimming periods are composed of longish swims a few metres below the surface with a consistent heading, interspersed with a brief return to the surface and reorientation [44].

Specifically, for an individual *i* at position x_*i*_(*t*) and time *t*, travelling with active velocity v_*i*_(*t*) = *s*(cos *α_i_*(*t*), sin *α_i_*(*t*)) where angle *α_i_*(*t*) denotes the active heading, then at time *t* + *t* (where *t* is small) we have:

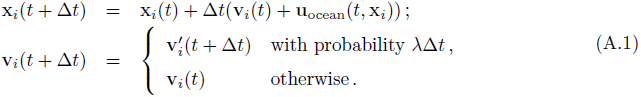

where 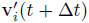 is the new velocity chosen at time *t* + *t* if a reorientation has occurred. The critical statistical inputs and parameters necessary to describe this process are as follows:

1. the average speed, denoted *s*;
2. the average rate of reorientations, λ
3. the directional distribution *q*(α) for turning into a new heading denoted by angle *α* ∈ [0, 2*π*).

It is via the directional distribution that navigating cues can be entered, by biasing turns into specific angles. As described in the main text, we employ the von Mises distribution, a *de facto* standard for directional datasets and widely adopted in theoretical and experimental studies of animal navigation (e.g. see [4]).

### Appendix A.2. Fully continuous model (FCM)

The scaling process that allows us to derive the FCM has been described previously (see [22, 46]). Defining *p*(x, *t*) to be the turtle population density (turtles per km^2^) at position x and time *t*, the dynamical change in *p* is governed by

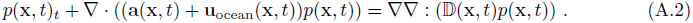

The above takes the form of a advection-anisotropic di usion equation, where advection is decomposed into components due to oriented swimming by the turtles (a(x, *t*)) and drift due to ocean currents (uocean). The anisotropic di usion, with di usion tensor D(x, *t*), emerges from uncertainty in the navigational choice. Notably, a(x, *t*) and D(x, *t*) are calculated directly from the inputs to the ABM and, under the von Mises distribution (1), can be explicitly calculated [23] as:

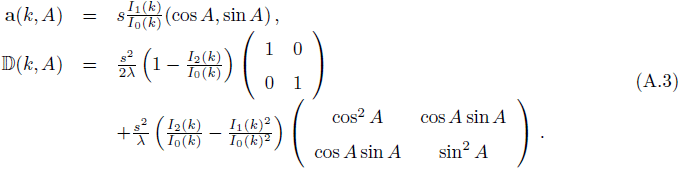

As *k* becomes large, D tends to the zero matrix and a(*k, A*) ✒ *s*(cos *A*, sin *A*). This corresponds to a “perfect navigator” that is able to move exactly according to the dominant direction. On the other hand, as *k* tends to zero a(*k, A*) ✒ 0 and 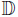 converges to an isotropic di usion tensor, corresponding to random movement. Note that for a relatively weak navigating field the model (A.2) can be further simplified, see [27], however we solve the full equation in the present paper.

## Appendix B. Navigating cues and fields

The consolidated navigating field, w_C_(x, *t*), combines information from the four individual navigating fields, w_T_, w_I_, w_O_, w_W_.

### Appendix B.1. Navigating fields for magnetic cues, w_*T*_ and w_*I*_

For w_T_ and w_I_ we assume turtles have an innate ability to detect and recall geomagnetic field information. Taking field intensity as an example, we assume its value is imprinted at the island and, following its subsequent displacement, a turtle experiences a bias according to the size and gradient of intensity di erence between that of its current position and that of the island.

Mathematically, we specify *M*_T_(x) as the magnetic intensity at position x and set

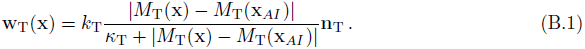

In the above n_T_ is the unit-length vector that points in the direction of the gradient of intensity di erence between the individual’s current position and that of Ascension Island. The above form stipulates that the strength of response to intensity di erences increases from negligible values close to the island, where intensity di erences are small, to a maximum value *k*_T_ when far from the island, presuming saturation in the detection mechanism. The sensitivity parameter _T_ reflects the subtlety to which intensity di erences can be detected.

Responses to magnetic inclination are treated similarly, where we set

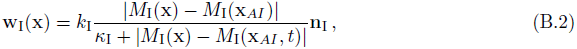

with equivalent definitions for the various parameters.

It is noted that we currently ignore the impact of temporal variation (diurnal, secular) or spatially localised anomalies on the size of field properties: *M*_T_(x) and *M*_I_(x) are the mean values generated by a standard geomagnetic field model across the study region, for the specific release times used in simulations.

### Appendix B.2. Navigating fields for odour plumes, w_*O*_ and w_*W*_

Fields w_O_ and w_W_ describe navigational responses to oceanborne and windborne plumes of some substance generated at the island. Simplistically, odour plume responses in animal navigation follow a pattern of chemoreception followed by movement upflow (see [61]), the latter termed anemotaxis for air currents and rheotaxis for water currents. This combination has been well documented in many species, including aquatic species such as sharks [24, 17] and lampreys [26]; we remark again that careful analyses of turtle tracks indicates a possible rheotactic response in marine turtles [28].

To model this behaviour in relatively simple terms, we take

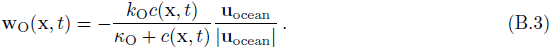

to describe response to an ocean-borne plume. In the above equation *c*(x, *t*) defines the substance concentration, the dynamics of which are defined below, while *k*_O_ and _O_ respectively define the strength and sensitivity of response to the chemical signal. The component u_ocean_/ |u_ocean_| describes the unit length vector in the direction of the local current, and the negative sign reflects that movement is upflow (i.e. against this direction). An equivalent formation is taken for wind-borne plumes,

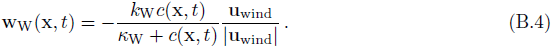

Each of the above navigational responses depend on a substance concentration, *c*(x, *t*), the dynamics of which are described by a standard advection-di usion-reaction model as follows:

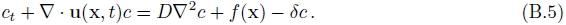

Convection arises from transport by wind or ocean currents, determined by velocity vector fields u(x, *t*) = u_wind_ or uocean respectively: Appendix D.1 details the public datasets used. *D* is a di usion coe cient, although it is noted to be relatively small compared to advection. The function *f*(x) describes production/release from the island; these could range from scents produced by island plants to coastal mud. We model this production via a two-dimensional Gaussian function centred at x_AI_,

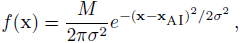

where *M* denotes the rate at which the substance is released and the parameter *σ* is taken to be su ciently small that any emission is effectively confined to the island. The decay coe cient δ = ln 2/*τ* accounts for subsequent “decay” of the substance, although this could also describe consumption by marine organisms or movement out of the relevant layer of detection; we note that we define this with respect to the standard notion of its half-life, *τ*.

## Appendix C. Model initial conditions

In the ABM each turtle is given a random starting location such that its distance from the island is distributed normally (with mean distance 300 km and standard deviation 10 km) and its angular position taken from a uniform distribution between 0 and 360°. The initial velocity is selected from the von Mises distribution (1) according to the navigating information provided from its starting location. Initial conditions for the macroscopic model are selected as

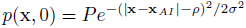

where *ρ* = 300 km, *σ* = 10 km and *P* is chosen to generate a density commensurate with an initial population of size 100.

## Appendix D. Datasets and parameters

### Appendix D.1. Datasets

Velocity fields for ocean currents (uocean) are generated from HYCOM (the global HYbrid Coordinate Ocean Model, [8]), an ocean forecasting model forced by wind speed, heat flux and numerous other factors that has been subsequently assimilated with field measurements (from satellites, floats, moored buoys etc) to generate post-validated output. HYCOM data is recorded at spatial and temporal resolutions of 1/12° and day to day, hence is capable of reproducing both large scale persistent currents and localised phenomena such as eddies. We assume turtles remain mainly at the same depth, and restrict to the upper most later recorded by HYCOM (*z* = 0). HYCOM data used in the current study was downloaded from http://pdrc.soest.hawaii.edu/data/data.php. Velocity fields for ocean surface winds (uocean) are generated and interpolated from Advanced SCATterometer (ASCAT) measurements. The ASCAT satellite tracks the modified radar backscattering resulting from small scale disturbances at the ocean surface due to surface winds. To provide day-to-day and regularly-gridded (1/4°) data, ASCAT observations are combined with European Centre for Medium Weather Forecasts and validated against measurements from, for example, moored buoys [6]. Datasets for these studies have been downloaded from the same repository as for HYCOM.

Magnetic field data (*M*_T,I_) for the region surrounding the island was obtained through the online calculator provided by the National Oceanic and Atmospheric Administration (http://www.ngdc.noaa.gov/geomag-web/) using the International Geomagnetic Reference Field (IGRF) model (12th generation), [60]. This model o ers a standard mathematical description of the Earth’s main magnetic field and its secular variation, derived and validated by an international consortium of modellers, institutes and observatories involved in collecting magnetic field data. As remarked on above, for the present study we consider only the mean values for two key field properties (total intensity and inclination angle) provided by this model, over the study region and restricted to the single time points corresponding to release dates.

We remark that for ease of computation, all datasets are interpolated from their native resolutions and saved at the same regular lattice of intergrid point spacing 1/12°. HYCOM/ASCAT data is saved at the day to day resolution available in the public datasets, while IGRF data is saved for each of the dates corresponding to one of the release times.

**Table D.2:**
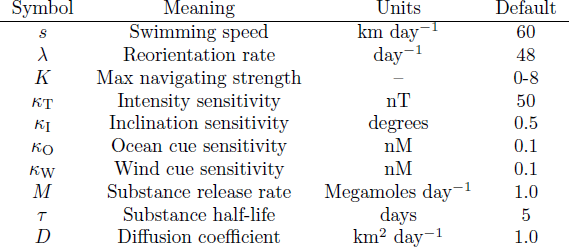
Model parameters and default values/ranges.

### Appendix D.2. Model parameters

We fix a default model parameter sets and ranges. Mean swimming speed is set at *s* = 60 km/day, below the upper limit of energetically sustainable swimming and in line with estimates based on tracking studies (e.g. [37]). The rate of reorientations is somewhat harder to gauge, and we propose a value according to two observations. First, analysis of turtle tracks during internesting periods indicate behaviours ranging from presumed active (and directed) swimming to feeding and resting [19, 55]: hence, it is reasonable to suggest that in general turtles do not navigate back to the island on a “24/7” basis. Second, the previously described studies of [44] suggests turtles surface and reorient approximately 3-4 times an hour during periods of active navigation. Combined, we propose an average daily rate corresponding to twice per hour for λ. It is noted, however, that perturbations to this value do not substantially alter results.

Given the unknown nature of any detected odours, associated parameters can only be stated in arbitrary fashion at present. In [29] the (horizontal) di usion coe cient was gauged to be of the order 10^2^ − 10^3^ km^2^/day: numerous orders of magnitude above typical molecular di usion coe cients (around 10^−9^ km^2^/day) to account for the turbulence-induced mixing by ocean currents. For our model, such mixing is explicitly captured in the spatio-temporally varying currents, and we therefore select a smaller value (*D* = 1km^2^/day) to represent fine-scale mixing below the resolution used in ocean current datasets. We note, however, that the contribution of di usive transport is negligible with respect to ocean transport: order of magnitude increases/decreases in *D* have no notable impact. No known data exists for release rates, which we arbitrarily set at 10^6^ moles per day: this is certainly not implausible for an entire island (for example, the earlier study of [29] employed a “modest guess” of 1 mole per second) and we take its half-life to be 5 days. Following the reasoning in [29] and taking the substance to become uniformly distributed in a 50 metre layer above (for air-borne) or below (for ocean-borne) the surface (see [29]), the above values generate *nM*–range concentration levels at distances extending 100s of kilometres from the island, plausible values for detection. We note that perturbations to the release rate simply act to modify the subsequent intensity of the signal: for example, scaling the release rate by a factor of 10 scale substance concentrations by a factor of 10.

Parameters associated with navigational fields will form the key focus for our investigation, and hence subject to variation. We note that navigational strengths are capped such that the absolute maximum strength varies in the range *K* ∈ [0, 8]. Any subsequent values of *k*(*t*, x) close to the upper limit would define an extremely precise navigator: for example, a value of *k* = 5 in equation (1) would generate turns within ±30° of the target angle *A* about 75% of the time. For comparison, values of *k* based on the swimming orientation of juvenile green and loggerhead turtles under laboratory-controlled conditions [2, 33] generate values in the range 1 − 2.5.

Sensitivity thresholds to magnetic intensity or inclination di erences are, naturally, di cult to gauge in animals. Studies in birds [57], fish [63] and bees [62] suggest that responses to intensity di erences are of the order of tens of nanoTeslas, while sensitivity to inclination angle di erences may extend down to fractions of a degree [48], see also [16, 66]. We therefore set κ_T_ = 50 *nT* and κ_I_ = 0.5°, although we will also explore the effect of perturbing from these reference values. It is noted that with these values, the strength of navigation response to the magnetic field decreases markedly below ~ 50 m from the goal, consistent with assumed ranges of an effective magnetic map [43].

Values for the sensitivity to air/waterborne substances is, naturally, somewhat arbitrary given their unknown nature. We note simply that the turtle olfactory system is highly developed (e.g. [3]) and, as an instance, responses to “naturally occurring” concentrations (pM range) of dimethyl sulphide (DMS) can be inferred from studies in loggerheads [12]. Here we set _O_ = _W_ = 0.1 nM, although again investigations have explored the effect of perturbations about this value.

### Appendix D.3. ABM Measurements

For data collection, in the ABM each turtle’s position is recorded once every 12 hours from release until it has either reached home or the tracking study has ended (*T* = 100 days). We set 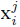 as the position of turtle *i* = 1 , . . . , *N* at time *j* = 0, 1/2, 1, . . . days and denote 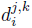 as the corresponding straight line distance between positions recorded at times *j* and *k*. We subsequently define the mean track distance (TD) as the population-mean length of turtle tracks,

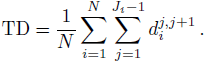

*J_i_* defines the number of recorded locations for turtle *i*. The mean straightness index (SI) measures the population-mean straightness of the turtle tracks,

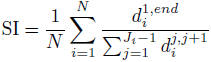

where 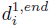 measures the distance between the first and last recorded position for turtle *i* (for example see [47]).

## Appendix E. Numerical methods

Numerical methods are adapted from our previous studies (e.g. see [46]) and described briefly below. Initially we note that the computational domain di ers from the study region, extending two degrees longer in each dimension: in effect, turtles can leave the 10° × 10° study region at one point and re-enter elsewhere. This practice allows specification of appropriate boundary conditions on the study region to be circumvented: on the computational domain edges we impose lossless boundary conditions for the turtles (turtles are, in effect, reflected) while chemical substances are allowed to flow across its edges. We note, however that the size of the computational domain is su ciently large that the exact choice has negligible impact on the results presented. As a second remark, we note that computer simulations run for *t* ∈ [−30, 100], where *t* = 0 denotes the point of release and *t* = 100 marks the final day of turtle tracking. The initiation of simulations 30 days before the release date ensures any odour plume has become established by the time of release.

### Appendix E.1. Hybrid ABM

The computational method used for the ABM combines stochastic simulations of each individual’s velocity-jump random path with (when necessary) a numerical approximation for the continuous equation for plume dynamics, equation (B.5). The numerical scheme is programmed using MATLAB. For the stochastic component, each particle is initiated at *t* = 0 with a position reflecting the stated initial conditions. Its movement path is subsequently generated through direct stochastic simulation of (A.1) according to the active and passive movement characteristics.

The time discretisation *t* used in simulation is suitably small, in the sense that a set of representative simulations conducted with smaller timesteps generate near identical results. For the selection of new active headings via the directional distribution given in (1) we employ code (circ vmrnd.m) from the circular statistics toolbox [7]. Currents and the inputs required for the active heading choice are interpolated from the native spatial/temporal resolutions in the saved variables to the individual particle’s continuous position x and time *t* via a simple linear interpolation scheme.

The numerical simulation of (B.5) adopts a simple Method of Lines approach, first discretising in space (using a fixed lattice of space *x*) to create a large system of ordinary di erential equations which are subsequently integrated over time. Approximation of the di usion term adopts a simple (second order) central di erence scheme while the advective component is solved via a third-order upwinding scheme, augmented by a flux-limiting scheme to ensure positivity of solutions (e.g. see [25]). Numerical integration in time assumes a simple forward Euler method. Discretisation adopts the same spatial resolution available for the datasets (Δ*x* = 1/12°) and the same time step used in the stochastic simulations (Δ*t* = 1/1000 day), with the ocean current velocity vector field at computational time *t* obtained through linear interpolating from its day to day value in the public datasets.

### Appendix E.2. FCM

For the FCM we adopt the same scheme used to approximate chemical plume dynamics and augment it with a similar method for the advection/anisotropic-di usion model for *p*(x, *t*), equation (A.2). Indeed, the only significant di erence emerges in the “fully anisotropic” di usion term, which can be expanded into an advective and standard anisotropic-di usion component. This ad-vective component, along with advection terms arising from ocean currents and active directional swimming, is solved using third-order upwinding with flux-limiting, as described above.

The choice of finite-di erence discretisation for the anisotropic di usion terms is more specific: naive discretisations can lead to numerical instability for su ciently anisotropic scenarios (high *k* values). The method of [65] allows greater flexibility in the choice of *k*: in this scheme, finite di erence derivatives are calculated and combined along distinct axial directions: the axes of the discretisation lattice and the major and minor axes of the ellipse corresponding to the anisotropic di usion tensor. Under the moderate levels of anisotropy encountered here (restricted by the size of *K*) we obtain a stable scheme. For the time integration we invoke a simple forward Euler method with a suitably small time-step, chosen to be the same as that employed for the hybrid model to facilitate comparison. To verify the numerical method, simulations have been performed for smaller time steps and smaller *x*. The validity of the methods for solving the ABM and macroscopic model is further substantiated by the quantitative comparisons shown below.

## Acknowledgements.

We thank Thomas Hillen and Jonathan Sherratt for insightful comments on earlier drafts and the anonymous reviewers whose suggestions have improved the manuscript. KJP acknowledges the Politecnico di Torino for a Visiting Professorship position (2016-2017). AP was supported by MIGSAA (a centre for Doctoral Training funded by EPSRC (grant EP/L016508/01), SFC UK, Heriot-Watt University and the University of Edinburgh).

